# Invariant categorical color regions across illuminant change coincide with focal colors

**DOI:** 10.1101/2022.06.05.494863

**Authors:** Takuma Morimoto, Yasuki Yamauchi, Keiji Uchikawa

## Abstract

Are there regions in a color space where color categories are invariant across illuminant changes? If so, what characteristics make them more stable than other regions? To address these questions, we asked observers to give a color name to 424 colored surfaces, presented one at a time, under various chromatic illuminants. Results showed a high degree of categorical color constancy, especially under illuminants that occur in the natural environment. It was also shown that surfaces selected as a focal color (the best example of a color category) are more resistant to illuminant change than non-focal color samples. This might imply that categorically invariant regions might have become focal colors to facilitate object identification and communication with others under a variety of lighting environments. We additionally ran an asymmetric color matching experiment to quantify the shift of color appearance induced by illuminant changes using surfaces that were all named gray, thereby disentangling the appearance-based color constancy from the categorical color constancy (which are often confounded). Results suggested that the appearance of color samples largely shifted due to illuminant changes even though all samples were named gray; showing that the constancy of a color category is substantially more robust than the constancy of color appearance.

## 1. Introduction

Physical parameters underlying color variation are continuous, and the associated subjective experience also varies continuously from one color to another. Yet, at the same time color perception can be discrete: we can count how many colors there are in a rainbow. This discrete nature is known as categorical color perception, and it offers a prime example to use colors in a daily life. Importantly, our visual system flexibly uses such a continuous-discrete duality to achieve diverse visual tasks from discriminating subtle color differences to effectively communicating with others using color (e.g. can you pass me the red coat?).

The interdisciplinary nature of color categories has naturally attracted a wide range of fields including biology, psychology, and linguistics. In the long history of research, many critical questions have been raised, investigated, and discussed. Are there ‘basic’ color categories in individual cultures (Berlin & Kay, 1969)? How are they different or shared across cultures (Uchikawa & Boynton, 1987), and are there universal constraints on the formation of color categories (World Color Survey, Kay & Cook 2016; Kay & Regier 2003; Kay & Regier 2007; Lindsey & Brown 2006; Lindsey & Brown 2009)? Do the linguistic labels we give to colored stimuli affect their color appearance (Sapir-Whorf hypothesis, Kay & Kempton, 1984)? Is it necessary to have linguistic labels to perform color categorization (Siuda-Krzywicka et al. 2019)? Does the strength of categorical effects differ between the right and the left visual field (Gilbert et al. 2006; Drivonikou et al. 2007; Gilbert et al. 2008)? Moreover, many psychophysical studies have demonstrated functional benefits to having color categories (Witzel & Gegenfurtner, 2018; Witzel, 2019). One representative example is the reduction of response time in a supra-threshold discrimination task when discriminating between colors in different color categories compared to the discrimination of colors within the same category (Bornstein & Korda, 1984). A cross-language study went further and showed that this categorical facilitation effect can be specific to the language that individuals use (Winawer et al., 2007). Curiously, certain color categories emerge only in certain conditions. The term ‘gold’ or ‘silver’ is used only for a glossy surface that exhibits specular reflection (Okazawa, Koida & Komatsu, 2011) or the term ‘brown’ is used only to describe the color of a surface, not the color of a light (Uchikawa, Uchikawa & Boynton 1989; Buck et al, 2016). Also color classification and categorization tasks are major interests in computer vision (Parraga & Akbarini, 2020). A recent study investigated the formation of color categories in an artificial visual system (de Vries et al. 2021), but their similarity to the human categorical system has not been formally evaluated.

These studies brought the whole picture of categorical color perception into sharp focus from wide perspectives. However, one crucial but missing element in these past studies is that in the real world, lighting environments largely change over space and time. For color categories to be useful in everyday tasks, one category should not easily change to another when a scene illuminant changes. Some studies have explicitly measured category-based color constancy. Uchikawa et al. (1989) might be the first study that introduced a color naming task to study color constancy, where the effect of chromatic adaptation was investigated. Troost and Weert (1991) replicated a ‘Mondrian’ experiment originally conducted by Arend and Reeves (1986) using both color naming and color matching tasks. They emphasized that color naming is potentially more suitable than color matching as a measure of color constancy because it is more naturalistic and therefore easier for observers to perform. Hansen et al. (2007) investigated the independent contributions of the spatial and temporal contexts on categorical color constancy. More recently a series of studies measured color naming patterns using color patches under simulated illuminant changes on a computer monitor (Olkkonnen, Hansen & Gegenfurtner, 2009) and Munsell color chips presented under real illuminations (Olkkonnen et al. 2010). It was shown that categorical color constancy holds slightly better for real surfaces than simulated surfaces, but more importantly both studies agreed that humans perform robust color categorization under different illuminants. Moreover, categorical response patterns were consistent across observers. One experiment conducted using a highly naturalistic experimental set-up also supported the robust categorical identification of colored papers under several lighting environments (Granzier, Brenner & Smeets, 2009). Most recently, Ma et al. (2018) used a three-channel LED and showed that categorical constancy holds generally better for red and green illuminants than for blue and yellow illuminants.

Basic aspects of categorical color constancy were uncovered by these efforts. Yet considering the large volumes of studies on color constancy (Foster, 2011), our knowledge on its categorical aspect is still largely limited.

We therefore conducted two types of psychophysical experiments to further investigate the mechanisms underpinning categorical color constancy, using real surfaces and real illuminants. Preliminary results and analysis of this study were presented at past research conferences (Uchikawa et al. 2002; Uchikawa et al. 2004), and this paper provides the full contents of the study. Aside from the potentially better constancy for real surfaces than for two-dimensional stimuli, the use of real stimuli has an advantage in that the stimulus range is not constrained by the color gamut of a monitor. We leveraged this strength and measured color naming patterns under a wider range of illuminants than past studies, including illuminants close to the spectrum locus. The first experiment was to name each of 424 color samples, presented one at the time on a gray surround placed under a colored test illuminant, using one of 11 basic color terms. We were particularly interested in (i) whether any regions in a color space remain invariant in their color category against illuminant changes and if so (ii) what makes those regions more stable than others. Such invariant colors could serve particularly important roles in visual tasks that might heavily rely on color categories. In a second experiment, we measured how much the color appearance of the surfaces named by the same color category shifts in response to illuminant color changes. In a hypothetical scenario where color appearance does not change at all thanks to the appearance-based color constancy, naturally, color categories should not change either. This second experiment was performed to disentangle two types of color constancy mechanisms which are often confounded in previous studies.

## 2. General Method

### 2.1 Test surface

All experiments used 424 color samples developed by the Optical Society of America (OSA, currently Optica), known as OSA uniform color scales (OSA-UCS). Each color sample has three coordinates j, g and L which stand for jaune (meaning yellow), green and lightness, respectively. A perceptual color difference between a color pair in OSA-UCS is defined as an Euclidean distance in j, g and √2L three-dimensional space (i.e. the L axis should be weighted by the factor of √2). Figure 1 shows the coordinates of all color samples. Each subpanel combines samples at two lightness levels (except for the plane of L = -7). At even lightness planes, the chromatic coordinates are all even integers by the design of the color system. In the same way at odd lightness planes, chromatic coordinates are always odd integers. For this reason there are no overlapping data points in Figure 1. We measured the spectral reflectance of each color sample using a spectrometer from 400 nm to 720 nm in 10 nm steps (X-Rite, Gretag Macbeth SpectroEye), and their CIE 1964 10-deg xy chromaticities under equal energy white illuminant are shown in the inserted horseshoe diagrams. More details of the color system are documented elsewhere (MacAdam, 1974; MacAdam, 1978), but the system was shown to be effective to capture color categorization patterns, where discrete and perceptually-uniform sampling of colors are key considerations (e.g. Boynton 1987; Boynton, Maclaury, & Uchikawa 1989; Boynton et al. 1989).

**Figure 1:**
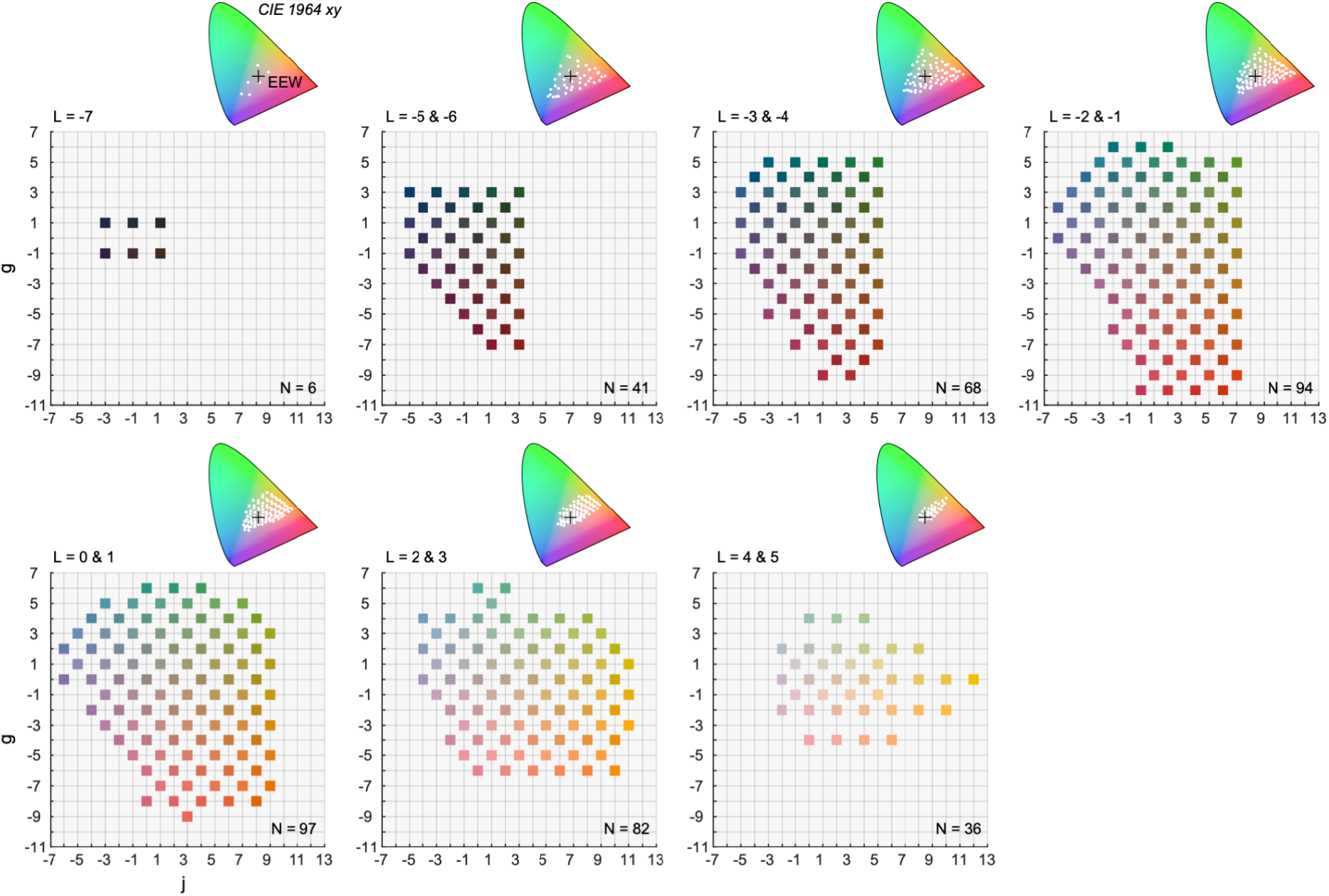
OSA uniform color system (*OSA-UCS*) used in all experiments. Each panel shows chromatic coordinates of color samples at different lightness levels. Inserted xy chromaticities were calculated under equal energy white illuminant.

### 2.2 Observers

Six observers participated in Experiment 1-1 (AH, KS, KU, MM, YE, and YY). Three of the six observers (KS, KU and YY) were further recruited for Experiment 1-2 and for Experiment 2. They were all screened to have normal color vision using Ishihara pseudoisochromatic plates. All observers were native Japanese speakers. Raw categorical responses were thus obtained in Japanese, which were translated to English for reporting purposes. The translation was straightforward as English speakers and Japanese speakers have largely equivalent basic color categories (Uchikawa & Boynton, 1987).

## 3. Experiment 1-1

### 3.1 Illuminant condition

For test illuminations, we used five illuminant spectra generated by a liquid crystal projector (SHARP LCP). Figure 2 (a) shows their spectral compositions together with the experimental set-up. The light emitted from the projector reached the test surface via the reflection of a mirror, and the light source was not directly visible to observers. Observers sat 53 cm from the test sample. For each trial, the experimenter carefully mounted a test color sample on a 5° × 5° square pedestal with a few mm height from a 50°×40° gray surrounding surface (lightness level equivalent to *OSA L* = -2). There were two ways to illuminate the test color sample. For (i) the whole illuminant condition as shown in panel (b), a color sample and the surrounding gray region were both illuminated by the test illuminant. This standard condition allowed us to measure the categorical patterns when categorical color constancy works to some extent. However, to quantify the degree of color constancy, we thought that it was desirable to know the categorical patterns without color constancy as a baseline measurement. This is a problem especially for color samples that are located around the center of a large color category because their category may not change even under large illuminant changes. In this sense, some colors have some baseline stability even without any color constancy mechanism. Thus, in addition to the standard whole-illuminant condition, we used (ii) the spot illuminant condition where only the test color sample was illuminated by the test illuminant while the surrounding region was illuminated by a 6,500K white light as depicted in panel (c), which was made possible by virtue of the use of an LC projector. This spot-illuminant condition fully eliminated cues to the color of test illuminant and created a condition where the color of the test illuminant appears to belong to the color of the test sample, thereby allowing us to measure color naming patterns under the test illuminant without color constancy. The naming patterns in the spot illuminant condition effectively defined the baseline for each color sample. The square pedestal was used to raise the color sample a few mm which minimized the amount of illuminant leaking into the surrounding region. We found this trick useful to make the test color sample appear as if the spectral reflectance of the surface changed when the test illuminant changed (i.e. no constancy).

**Figure 2:**
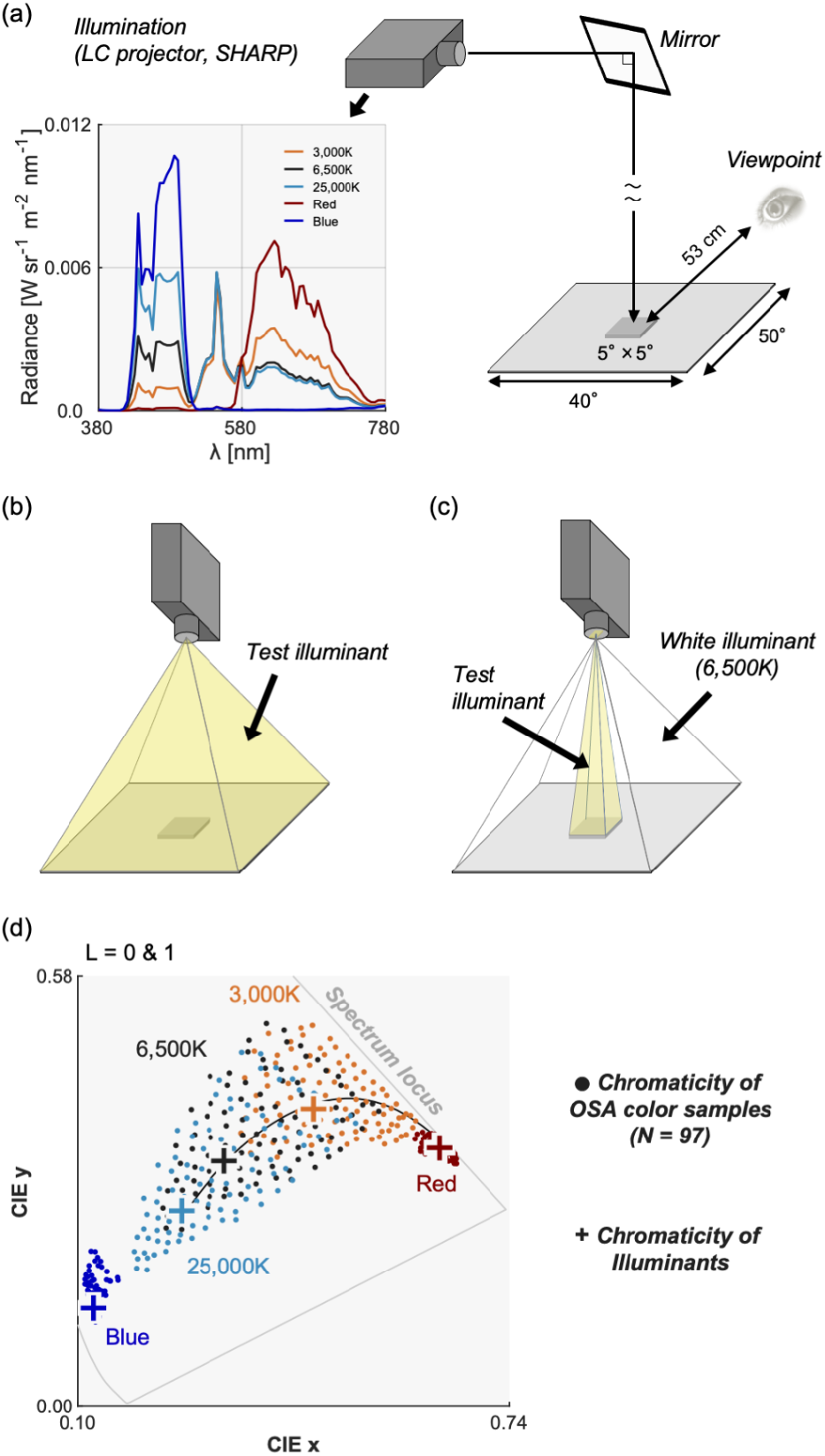
(a) Experimental set-up in Experiment 1-1. The 5°×5° test color sample was presented on a uniform gray surrounding. For the whole illuminant condition, both the test color sample and the surrounding gray region were illuminated while for the spot illuminant condition only the test sample was lit by the light source. (b) Whole illuminant condition. Test color sample and surrounding region are both illuminated by the test illuminant. (c) Spot illuminant condition. Only the test color sample is illuminated by the test illuminant and the surrounding region is illuminated by a 6,500K white light. (d) The influence of illuminant change on the CIE 1964 xy chromaticity of 97 color samples (*L* = 0 and 1). The plus symbols depict the chromaticities of test illuminants that varied along and beyond the black body locus shown by the black solid curve.

To show the magnitude of each illuminant change, Figure 2 (d) shows chromaticities of five test illuminants and the associated chromatic distribution of the 97 color samples at a mid lightness level (L = 0 & 1) under each illuminant. The color temperatures *3,000K* and *25,000K* were selected as they were around the edges of the natural daylight variation (Morimoto et al. 2022). The ‘Red’ and ‘Blue’ illuminants were achieved by activating only red or blue phosphors of the projector, respectively. Thus, they had particularly high stimulus purity, under which color samples had chromaticities tightly clustered near the spectrum locus. These illuminants were chosen to investigate the limit of categorical color constancy outside the natural illuminant variation.

### 3.2 Procedure

Observers named 424 color samples under 10 different illuminant conditions (5 illuminant colors × whole and spot conditions), using one of 11 basic color categories: red, green, yellow, blue, brown, orange, purple, pink, gray, white and black. One session thus consisted of 4,240 responses, and all observers completed 2 sessions. After the categorization of all 424 color samples under each illuminant condition, observers were additionally asked to select a focal color for each color category. There was no time limit in each trial. Observers first adapted to a test illumination for five minutes via the reflection from the gray surrounding region with no test color sample presented. Color samples were presented one at a time in a random order. The order of illuminant condition was also randomized.

### 3.3 Results and Discussion

#### 3.3.1 Color naming patterns in the OSA color system

Figure 3 shows the color naming results under the *6,500K-whole* illuminant condition for an observer AH. Each data point was colored by the color term used by the observer. A bi-colored data points show the color name used for session 1 (outer color) and for session 2 (inner color). Color samples selected as focal colors are additionally marked by a white cross (if consistent across sessions) or a white plus (if not). The color categories spread across the OSA color system, and it is shown that some color categories (e.g. green) were used more than others (e.g. red, gray). The black and white were not used by this observer. The response patterns were roughly consistent for the other 5 observers.

**Figure 3:**
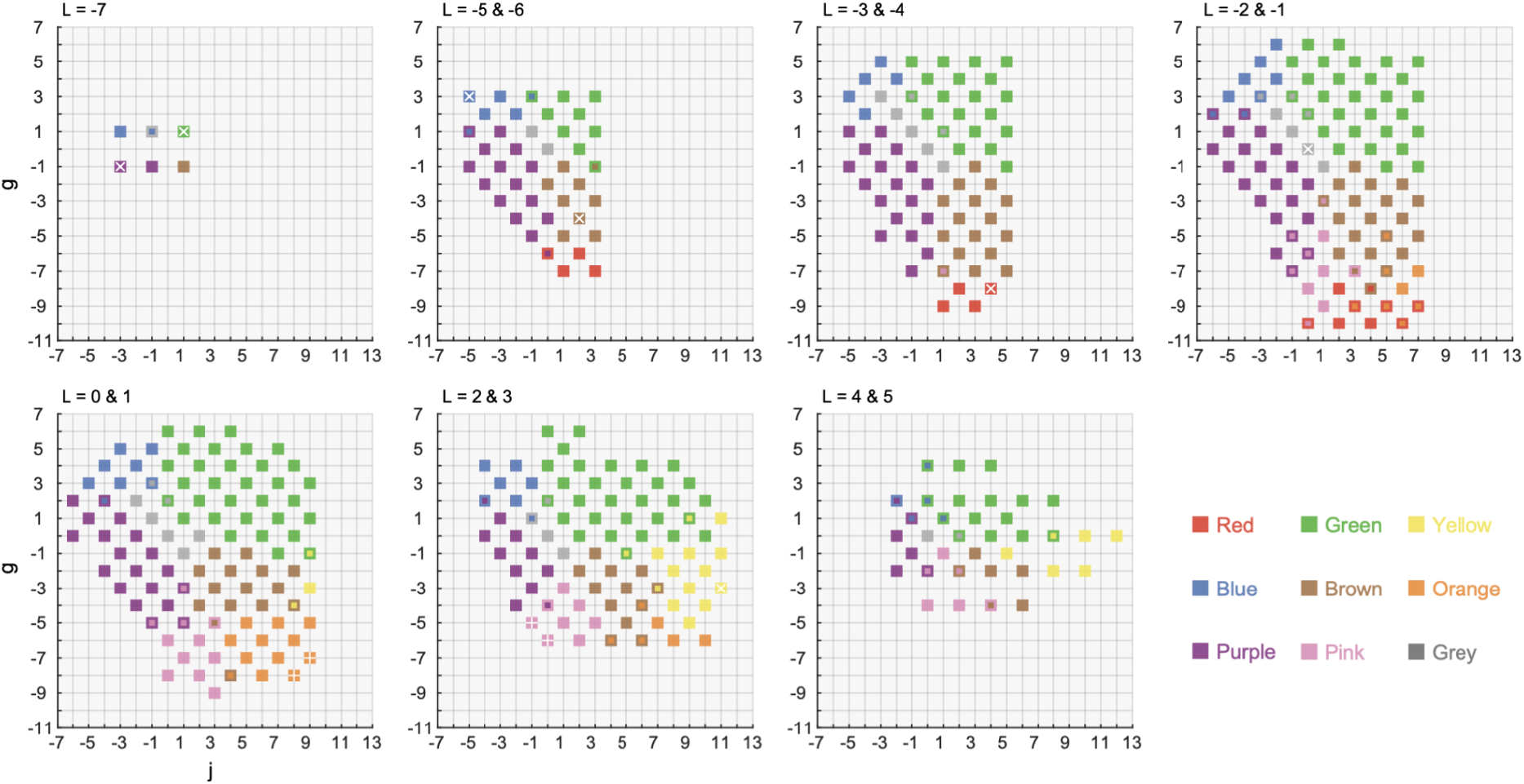
Color naming results under *6,500K-whole* illuminant for one observer AH. Data points marked by two colors show the color name used for the first session (outer color) and for the second session (inner color). The color sample chosen as the focal color is depicted by a white cross (if consistent across two sessions) or a white plus (if inconsistent).

Figure 4 (a) shows how color naming patterns change in response to illuminant change. For brevity, we show only the results for 97 color samples at L = 0 and 1. The upper subpanel re-plots the standard illuminant condition (*6,500K-whole*) for comparison. A perfectly color-constant observer would show the same categorical response pattern regardless of illuminant conditions.

**Figure 4:**
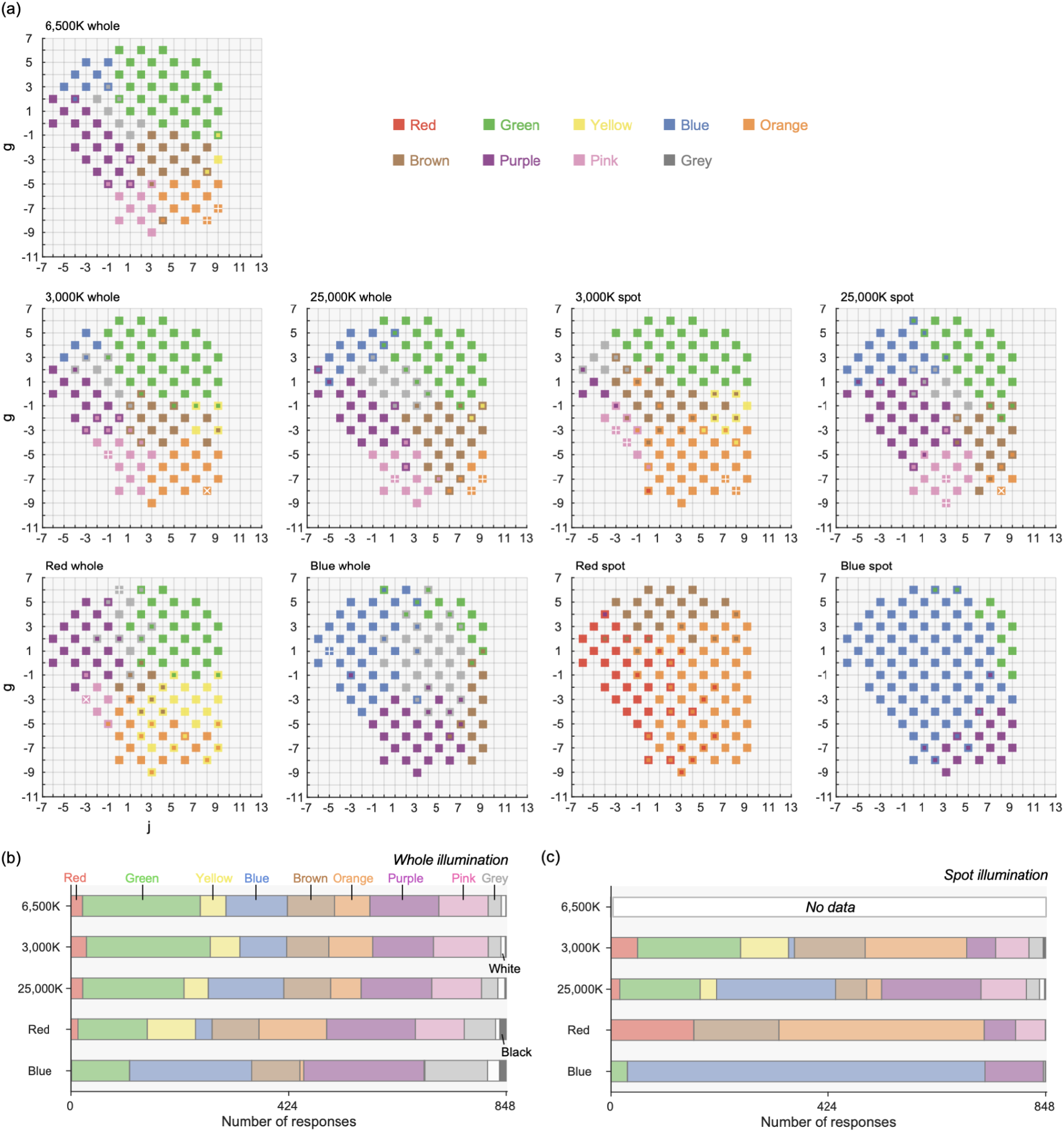
(a) Color naming results at L = 0 and 1 under different illuminant conditions for observer AH. The upper sub-panel replots the result for the standard illuminant condition for comparison purposes. The left and right four sub-panels depict results for the whole illuminant and spot illuminant conditions, respectively. Color samples that were selected as focal colors are marked by a white cross (if consistent across sessions) or a white plus (if inconsistent). (b, c) The number of categorical responses for the whole illuminant condition and spot illumination condition, respectively. The numbers were averaged across 6 observers.

For the *3,000K-whole* and *25,000K-whole* conditions, color categories are largely similar to those in the standard illuminant condition. In contrast, for the *Red-whole* illuminant, slightly more yellow and orange responses are seen around the lower part, and blue responses around the upper-left region were replaced by purple responses showing a bias towards the color of the test illuminants. Also the *Blue-whole* illuminant showed more prevalent blue and purple responses, showing a failure of categorical color constancy.

For the *3,000K-spot* and *25,000K-spot conditions*, though we still see a somewhat similar pattern to that of the standard illuminant condition, response patterns are more severely affected by the illuminant color than the whole illuminant conditions, as we had expected. Some color categories did not change, likely because the shift of chromaticity due to illuminant change was not large enough to alter categorical responses for those color samples, showing their baseline tolerances. For *Red-spot* and *Blue-spot* illuminants, however, response patterns are largely different, where mostly red and orange or blue and purple responses were recorded, respectively. Comparing these patterns to those in Red-whole and Blue-whole conditions allows us to appreciate how well categorical color constancy holds in their respective whole illuminant conditions.

Panels (b) and (c) summarize the number of color terms used for the whole illuminant and the spot illuminant conditions, respectively, visualizing how the relative frequency of categories varies across illuminants. There were 848 responses in total for each illuminant condition (424 color naming × 2 sessions), and the numbers were averaged across all 6 observers. The number of responses naturally varies across color categories. For example, red and yellow categories were used much less frequently than colors such as green, purple and pink. Also, panel (c) clearly visualizes that categorical patterns differ much from the whole illuminant condition, especially for the *Red* and *Blue* illuminants. The proportion of white and black responses were only 1.47% and 0.723% on average across observers, and thus we excluded these categories from subsequent analyses.

#### 3.3.2 Categorical centroids

We next quantified the magnitude of categorical shifts induced by the illuminant change. To define a representative location for each color category, we computed their centroids in the following way that is in accordance with past studies (e.g. Boynton, MacLaury & Uchikawa, 1989). For each observer, there were in total 848 responses, each labeled by a color naming. Then, for a specific color category, for instance red, we took all color samples named red, and calculated mean *j, g* and *L* values across the samples. The calculation was done separately for each illuminant condition. Figure 5 shows the centroids of color categories for each illuminant condition and the value in each circle shows the *L* value of the centroid. The size of each circle corresponds to the number of responses that went into the computation of the centroid. Then, for each color category, we calculated the OSA color difference *ΔUCS* between standard illuminant (i.e. *6,500K-whole*) and each illuminant condition using equation (1), where suffixes 1 and 2 indicate the standard illuminant and each illuminant, respectively. The weighting of square root two is applied to *L* values to account for the geometrical asymmetry in the color system (MacAdam, 1974).

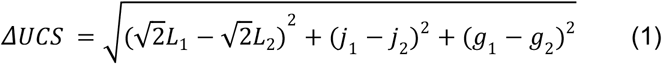

Figure 6 (a) shows *ΔUCS* values for whole and spot illuminant conditions, averaged across 6 observers. The absence of data points means that no observer used the color name for the condition. For data points where one or more observers showed no response, we calculated the average without those observers. In the *OSA-UCS*, neighboring colors have at least 2.0 *ΔUCS* (shown by horizontal magenta lines). We see that *ΔUCS* values in the whole illuminant condition are very small for (a) *3,000K* and (b) *25,000K* conditions, where values were lower than 1.0 in most categories and thus comfortably smaller than the minimum difference in the color system. As expected, *ΔUCS* values in spot illuminant condition were always larger than the whole illuminant condition.

**Figure 5:**
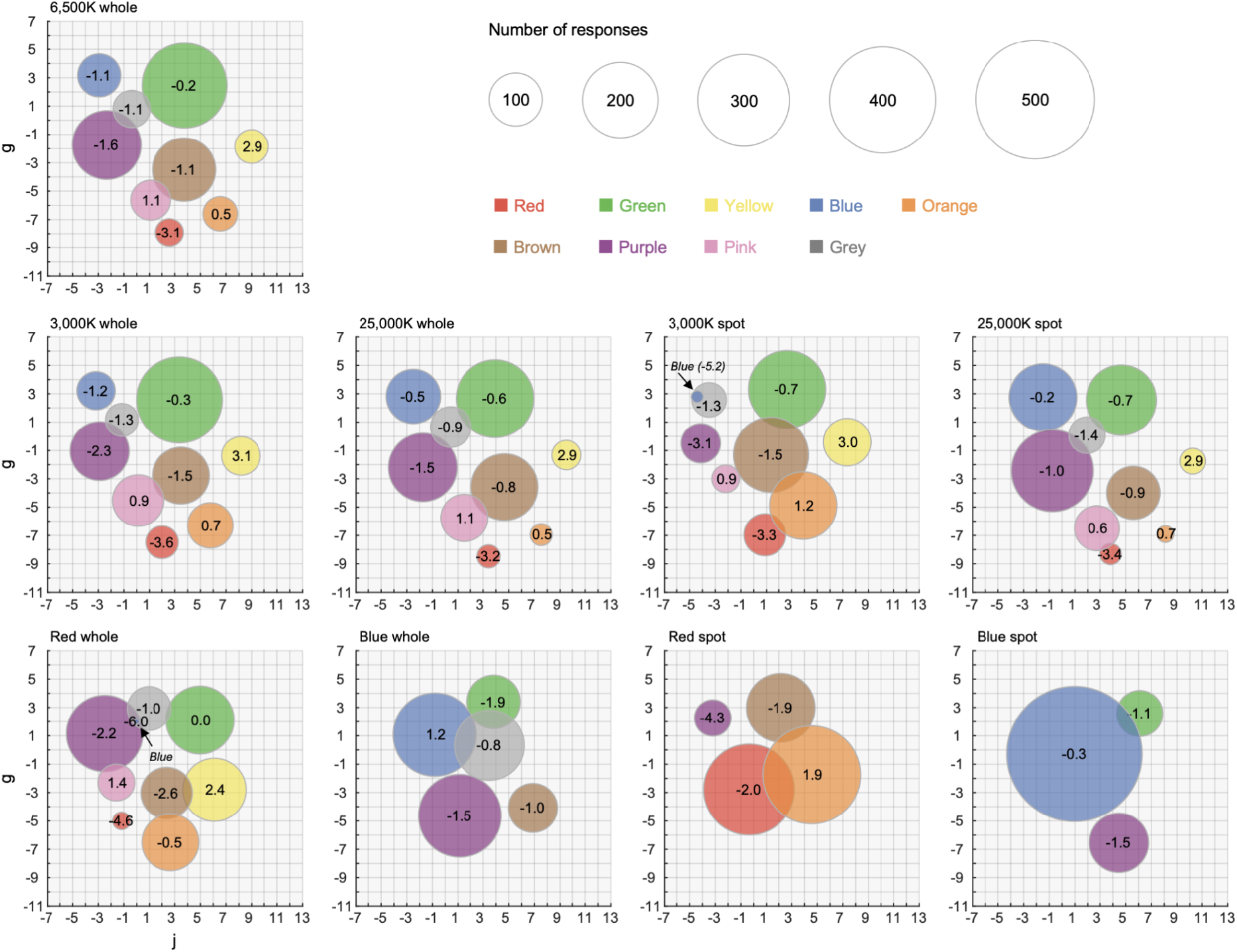
Centroids of color categories for observer AH. The size of each circle depicts the number of responses in the category. The value in each circle is the L value of each centroid.

**Figure 6:**
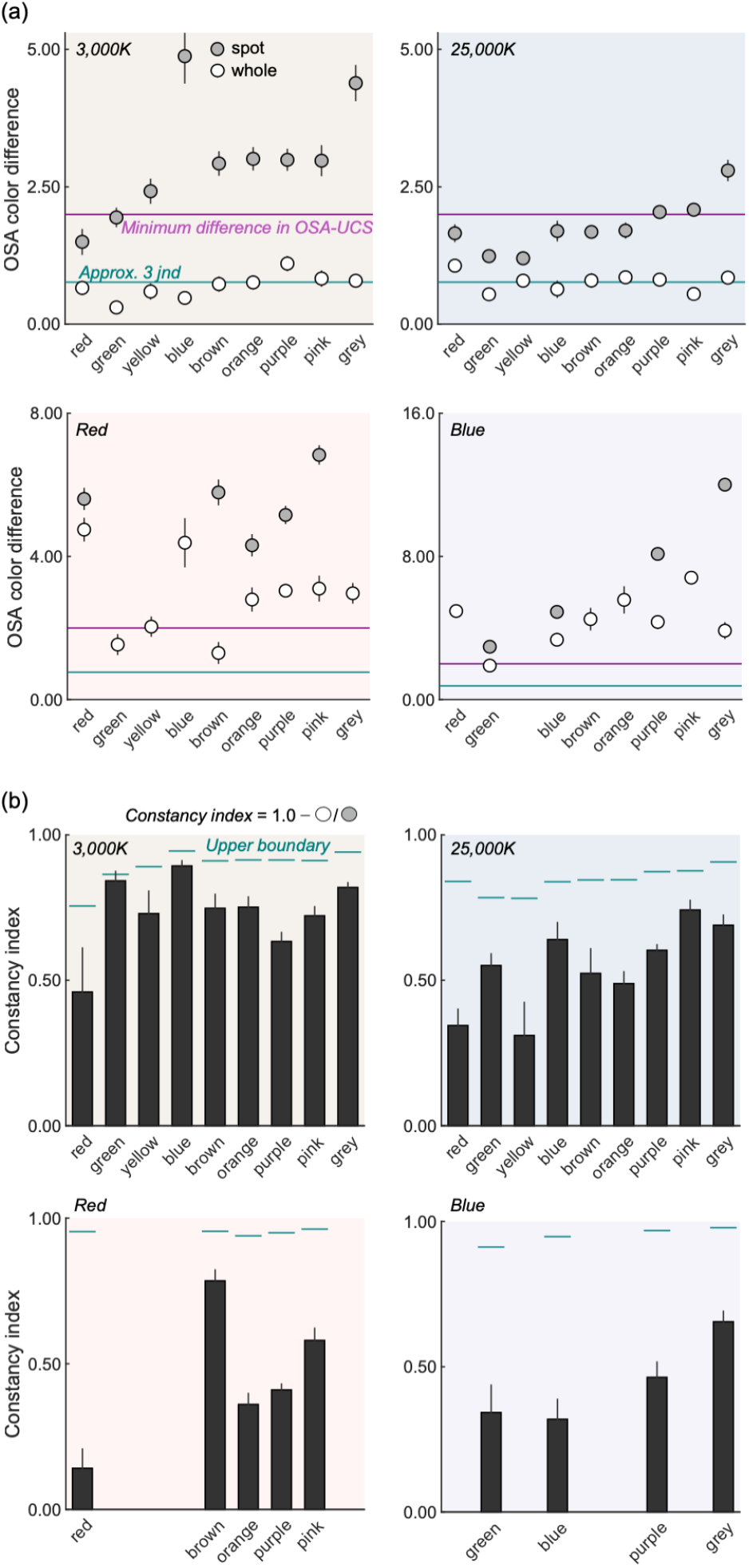
(a) The shift of categorical centroid in the *OSA* unit between the *6,500K*-*whole* illuminant condition and other illuminant conditions. The magenta horizontal line shows the minimum difference that can be defined in the *OSA-UCS*. The cyan horizontal lines indicate roughly 3 *jnd*s, estimated based on relation between *ΔUCS* and *DE2000* (see main text for details). (b) Constancy indices defined by the ratio of *ΔUCS* between the whole illuminant and the spot illuminant condition subtracted from one. The value one means perfect color constancy and zero shows no color constancy. The error bars show ± 1.0 S.E across observers in both panels.

Furthermore, to have a more concrete idea as to the magnitude of these shifts, we analyzed how *ΔUCS* values relate to another commonly used color difference metric *DE2000* defined in L*a*b* color space (Luo & Rigg, 2001). For this, we calculated *ΔUCS* values and *DE2000* values for all possible pairs of 424 *OSA* color samples (89,676 pairs). It was found that *ΔUCS* and DE2000 are highly linearly correlated (*r* = 0.906, p < 0.00001) and that 0.255 *ΔUCS* approximately corresponds to 1.0 *DE2000* which is often interpreted as one just noticeable difference (*jnd*). As shown by the cyan horizontal line in panel (a), 3 *jnds* were found to be roughly equal to the magnitude of shifts in the *3,000K* and *25,000K* whole illuminant conditions. However, it should be stressed that this relation between two color metrics is only a loose approximation and should not be taken as a generic conversion for other uses.

For the *Red* and *Blue* whole illuminant conditions, the color shifts are much larger than the *3,000K* and *25,000K* conditions, except for the yellow category in the *Blue* whole illuminant condition that was not used by any observer. And for spot illuminant conditions, some categories were used by no observers, shown by the absence of data points.

To summarize these in a more quantitative way, we next calculated constancy indices (*CI*) using equation (2). Here *a* and *b* show *ΔUCS* values in the whole illuminant condition and *ΔUCS* values in the spot illuminant condition, respectively. The value one and zero of the index means perfect and no constancy, respectively.

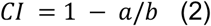

Panel (b) shows the indices for all illuminant conditions. Some categories were left blank because it was not possible to compute the index for these categories because of the absence of data points (as seen in panel (a)). It was shown that categorical constancy holds well for *3,000K* and *25,000K* though there are some areas of improvement, especially in the *25,000K* condition. For the *Red* and *Blue* conditions, color constancy was found to hold for some categories to some extent. It should also be noted that the blank data does not mean no color constancy. If anything they show high color constancy in a sense that no responses were recorded in the spot-illuminant condition but they emerged in the whole-illuminant condition thanks to color constancy. We also considered that the constancy index is likely limited by our absolute color sensitivity (*jnd*). Based on this thought we computed the upper-bound of constancy indices by taking the ratio between 0.255 *ΔUCS* (1.0 *jnd* as estimated above) and *ΔUCS* in the spot illuminant condition, which was then subtracted from one. This is shown by the cyan lines in panel (b). A constancy index between this line and 1.0 should be collectively interpreted as perfect constancy.

#### 3.3.3 Focal colors

Next we analyzed the focal colors, which might serve functional significance in mechanisms of categorical color perception because they are likely the first colors that come to our mind when we communicate using color terms. For the communication to be effective, however, the focal color needs to be shared across individuals at least to some extent and also needs to be robust across lighting environments. Based on these ideas, we have analyzed the locations of the focal colors to see (i) how they vary across individuals and (ii) how they change depending on the illuminant condition.

Here, to directly see what chromaticities were chosen as focal colors under each test illuminant (whole-illuminant) we plotted them in the CIE 1964 xy chromaticity diagram as shown in Figure 7 (a) - (e). The shape of symbols depicts observers. For each panel, the plotted *xy* ranges differ as shown in the diagram next to panel (a). Roughly speaking, we see trends that focal colors are clustered somewhat closely for the *6,500K, 3,000K* and *25,000K* conditions. For the *Red* and *Blue* illuminant conditions, the variation across observers seems slightly higher. However, it is curious that many color categories emerge even for these extreme *Red* and *Blue* illuminants which push all chromaticities towards very small regions in a color space close to the spectrum locus. In contrast, for spot illuminant conditions where no color constancy holds, as shown in Figure 5, only limited categories were recorded for *Red* and *Blue* illuminants, meaning that the emergence of various categories here shows the presence of color constancy.

**Figure 7:**
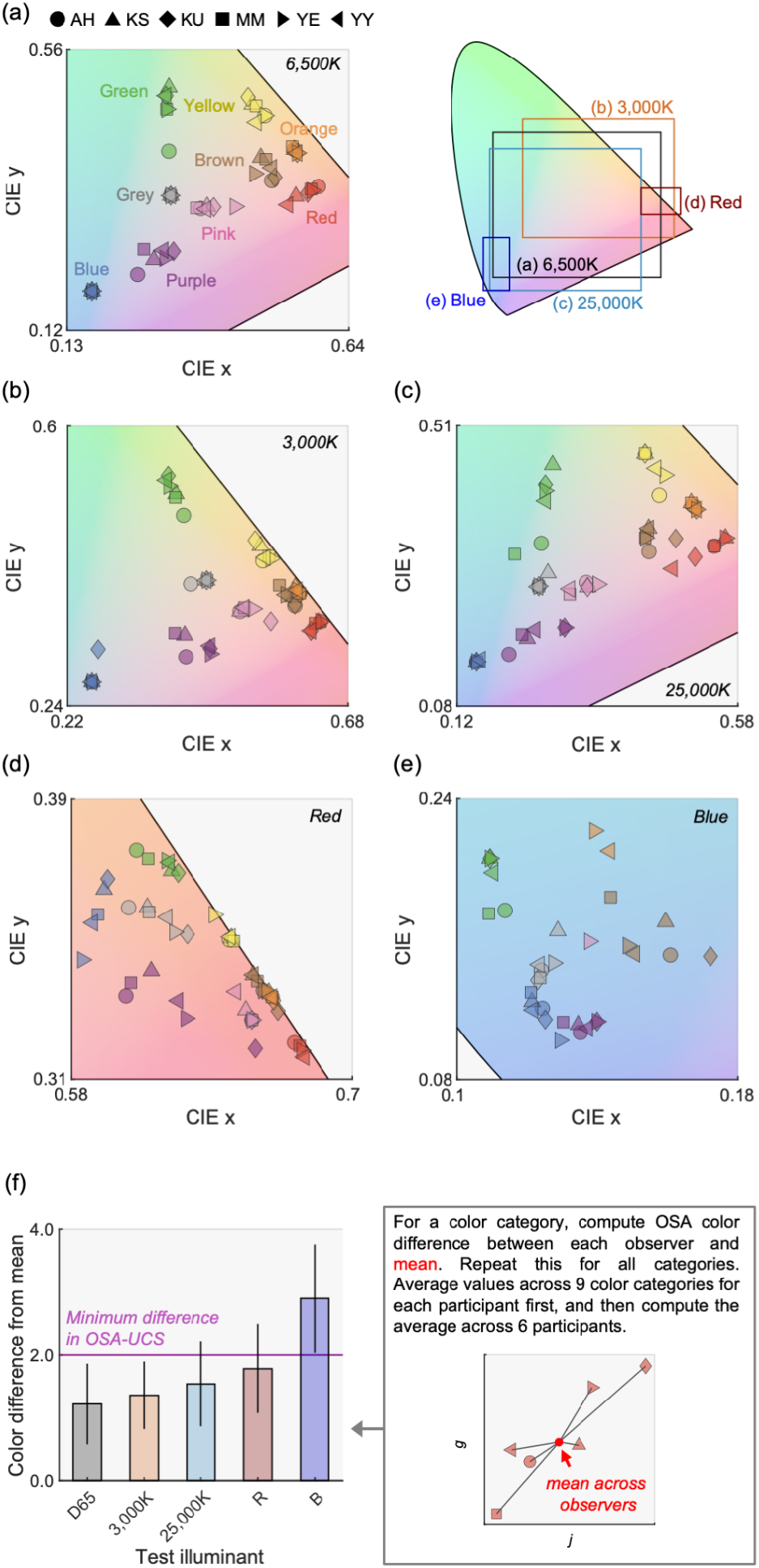
CIE 1964 xy chromaticity of color samples chosen as a focal color for (a) *6,500K*, (b) *3,000K*, (c) *25,000K*, (d) *Red* and (e) *Blue* illuminants (whole-illuminant condition). The color of the symbol depicts the color category and the shape shows observers. Each panel shows different *x* and *y* ranges, which are collectively shown in the inserted diagram next to panel (a). (f) Color difference in *OSA* unit between mean across observers and each observer. The value was averaged across color categories and then across observers. The magenta line shows *ΔUCS* = 2.0, corresponding to the minimum difference in the *OSA-UCS*. The error bars show ± 1.0 *S*.*E*. across color categories.

To quantify the variation of focal colors across observers, we calculated the mean *ΔUCS* between mean focal color point and focal color for each observer. The computation was done separately for each category first, and then took the average across categories. The right-hand part of panel (f) also describes this computation. We see that the variation across observers is smaller than the minimum difference in the color system for natural illuminants (*6,500K, 3,000K* and *25,000K*) and *Red* illuminant, suggesting that focal colors are relatively stable across observers for these illuminants. One-way repeated-measures ANOVA revealed that the main effect of illuminant conditions (*6,500K, 3,000K, 25,000K, Red* and *Blue*) is significant (*F(4, 20)* = 14.4, *p* = 0.000011). Multiple comparisons with Bonferroni’s correction reported the following relation: *6,500K, 3,000K, 25,000K* and *Red* < *Blue*. In other words, other than the *Blue* illuminant, the stability of focal color across observers was roughly equal.

These analyses generally supported the idea that focal colors were stable across at least the 6 tested observers. Next, we tested whether focal colors were stable across illuminants. For this, we first calculated the consistency of color category across illuminants for each color sample. For each color sample, we had 10 responses (5 illuminant colors × 2 sessions). Then, the consistency was defined as the proportion that the same color category was used for the color sample. For example, if the color sample was named by the same category 7 times out of 10 responses, this color sample had 70% consistency.

In Figure 8, this consistency is shown as the heatmap where darker blue indicates higher consistency. The colored symbols show the location of focal colors under *6,500K*, and the shape of symbols depicts observers. Roughly speaking it is shown that focal colors are almost absent around the color samples that have particularly low consistency. Instead they tend to fall on to color samples that have relatively high consistency though the exact trend depends on color categories. We see that this observation holds particularly well for categories such as green, purple, or brown.

**Figure 8:**
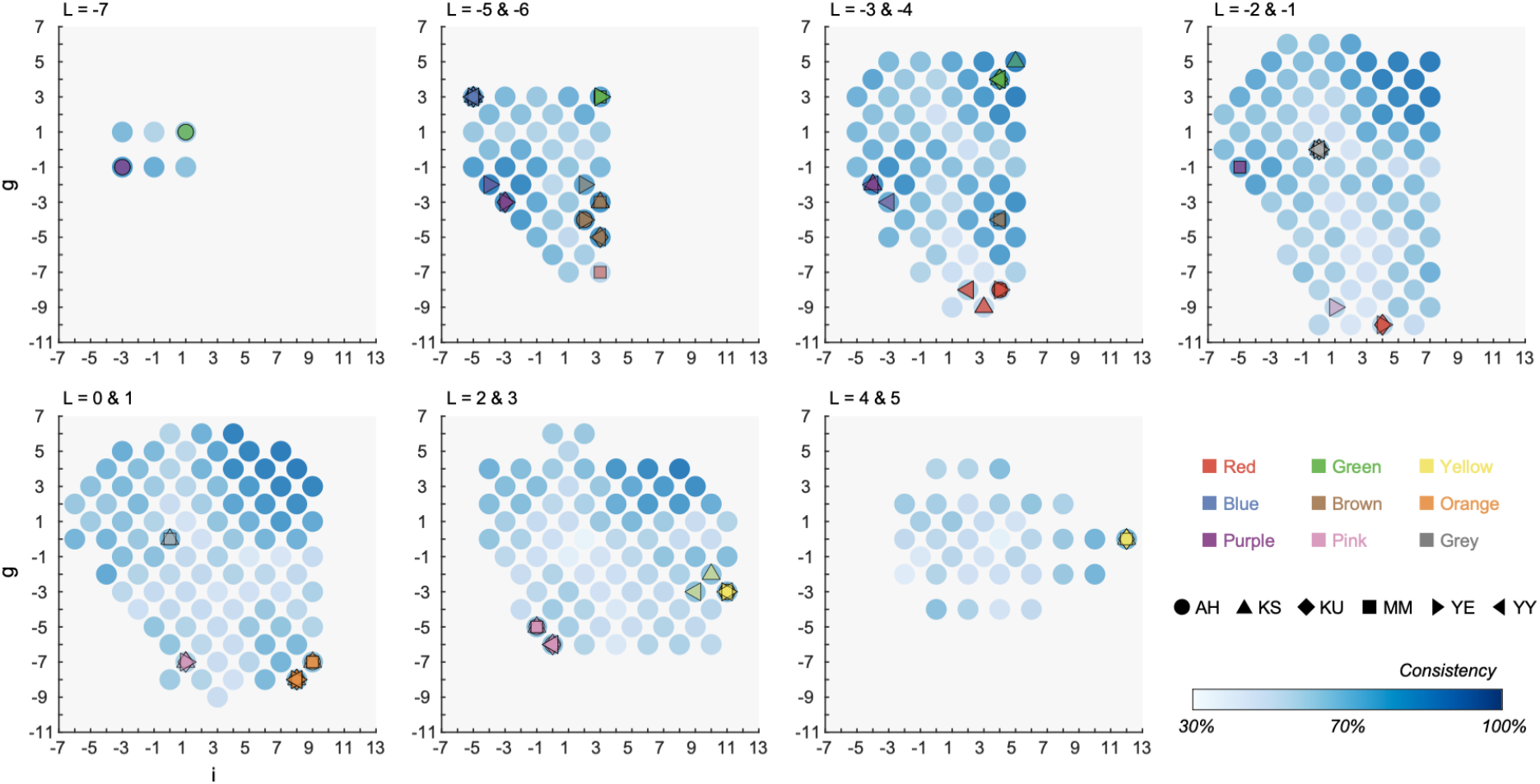
Consistency across 5 illuminants for focal colors and non-focal colors in the whole illuminant condition. The values were averaged across 6 observers. For each color sample, there were 10 responses (5 illuminant colors × 2 sessions). The consistency was defined by the proportion that a color sample was called by the same category. Colored symbols show focal colors selected in the *6,500K*-*whole* illuminant condition.

We next computed the average consistency separately for focal color samples and non-focal color samples. We first fixed a color category, and then we collected color samples that were selected as the focal color at least once under *6,500K*, and calculated mean consistency across the color samples. For non-focal colors, we needed to define which color sample belonged to which color category, as the color naming varies across illuminant conditions. Each color sample was classified to one color category based on the mode across 10 responses (i.e. color name that was used most frequently for the color sample).

Figure 9 (a) shows the consistency for focal color samples and non-focal color samples. A two-way repeated-measures ANOVA confirmed the significant main effect of the focality (*F*(1, 5) = 73.6, *p* = 0.000355, focal > non-focal) and the color category (*F*(8, 40) = 6.99, *p* < .00001). The interaction between the focality and the color category was significant (*F*(8, 40) = 2.85, *p* = 0.0132). Multiple comparisons by Bonferroni’s correction showed that the focal color had higher consistency than non-focal colors in most color categories as shown by asterisks in Fig. 9 (a). We next compared the consistency between color categories and found significant differences for the following categorical pairs: for focal colors (red < green, brown, purple; pink < brown, purple), non-focal colors (red < green, purple; green > pink, grey; purple > pink, grey). To summarize, these analyses suggested that focal colors are more stable across illuminants than non-focal colors, for most color categories.

**Figure 9:**
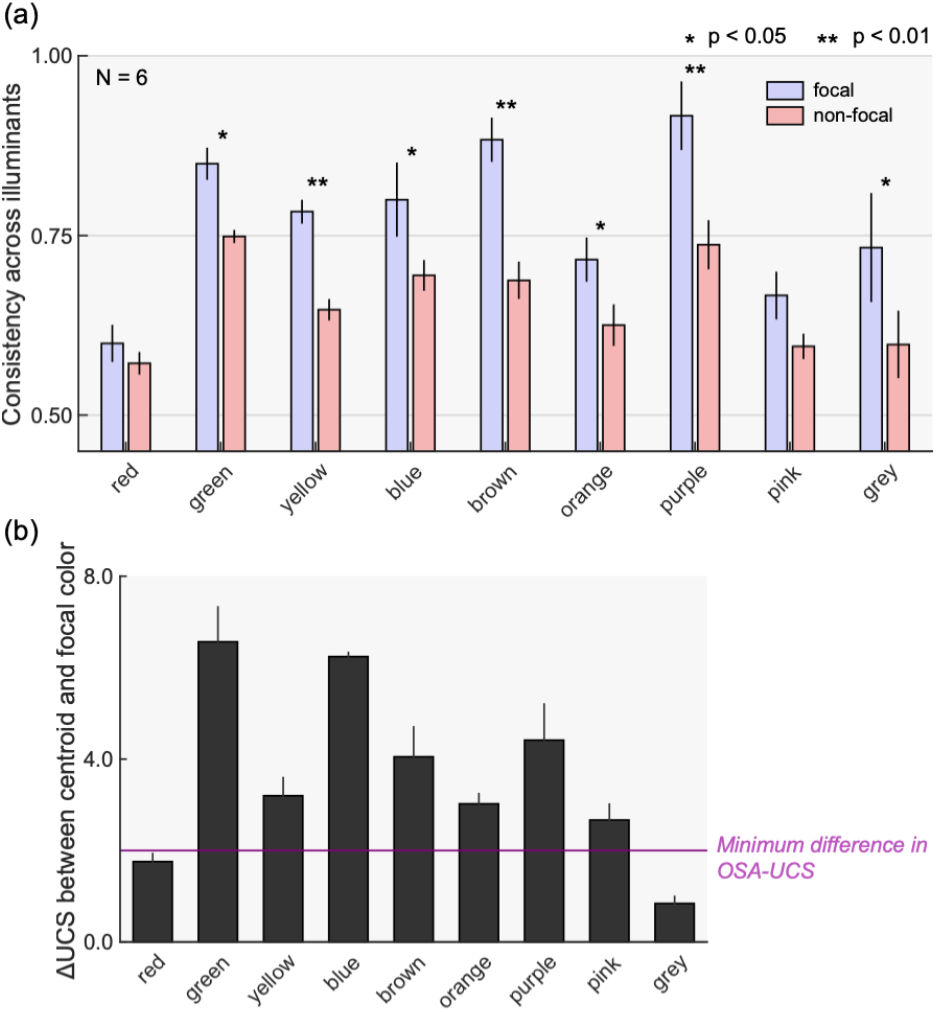
Consistency across illuminants for focal color samples that were selected at least once as a focal color in *6,500K* condition and non-focal color samples that were never selected as a focal color. The symbols * and ** indicates pairs where significant differences were found (Bonferroni’s correction applied). The error bars show ± 1.0 S.E across observers.

One remaining question is why focal colors are stable. One possibility would be that focal colors are simply located near the centroid of the color category, and thus illuminant changes are less likely to push them outside the category. To test this idea, we calculated *ΔUCS* between the centroid and focal color. The calculation was first done for each observer independently and averaged across two sessions. Then the *ΔUCS* values were averaged across observers. Rather surprisingly, we found that in most cases *ΔUCS* values are substantially higher than the minimum difference, indicating that focal colors do not necessarily coincide with categorical centroids. We see smaller *ΔUCS* values for red and gray categories, but the number of responses were lower in these categories. For these small color categories observers did not have an option to pick a color sample far from the categorical centroid as the focal color, which necessarily led to a small *ΔUCS* value here. Thus, these observations collectively reject the hypothesis that focal colors are stable merely because they are located around the center of color categories.

## 4. Experiment 1-2

### 4.1 Condition and procedure

One major finding in Experiment 1-1 was that invariant categorical regions coincide with focal color locations. However, the consistency values shown in Fig. 9 (a) directly depends on the number and variation of illuminants we used. We thus felt it would be desirable to validate this main finding using an expanded set of illuminants and decided to run a supplemental experiment. Figure 10 (a) and (b) show the spectra of seven illuminants used in this experiment. Two additional light sources were 1500K and violet, whose chromaticities were located between *3,000K* and *Red* and between *25,000K* and *Blue*, respectively. A different three-primary liquid crystal projector was used in this experiment (EPSON, EMP-54, spatial resolution 800 × 1300, horizontal scanning frequency 15 - 92KHz, vertical scanning frequency 50 - 85Hz). We used only the whole illuminant condition. Other experimental conditions and procedures were identical to those in Experiment 1-1.

**Figure 10:**
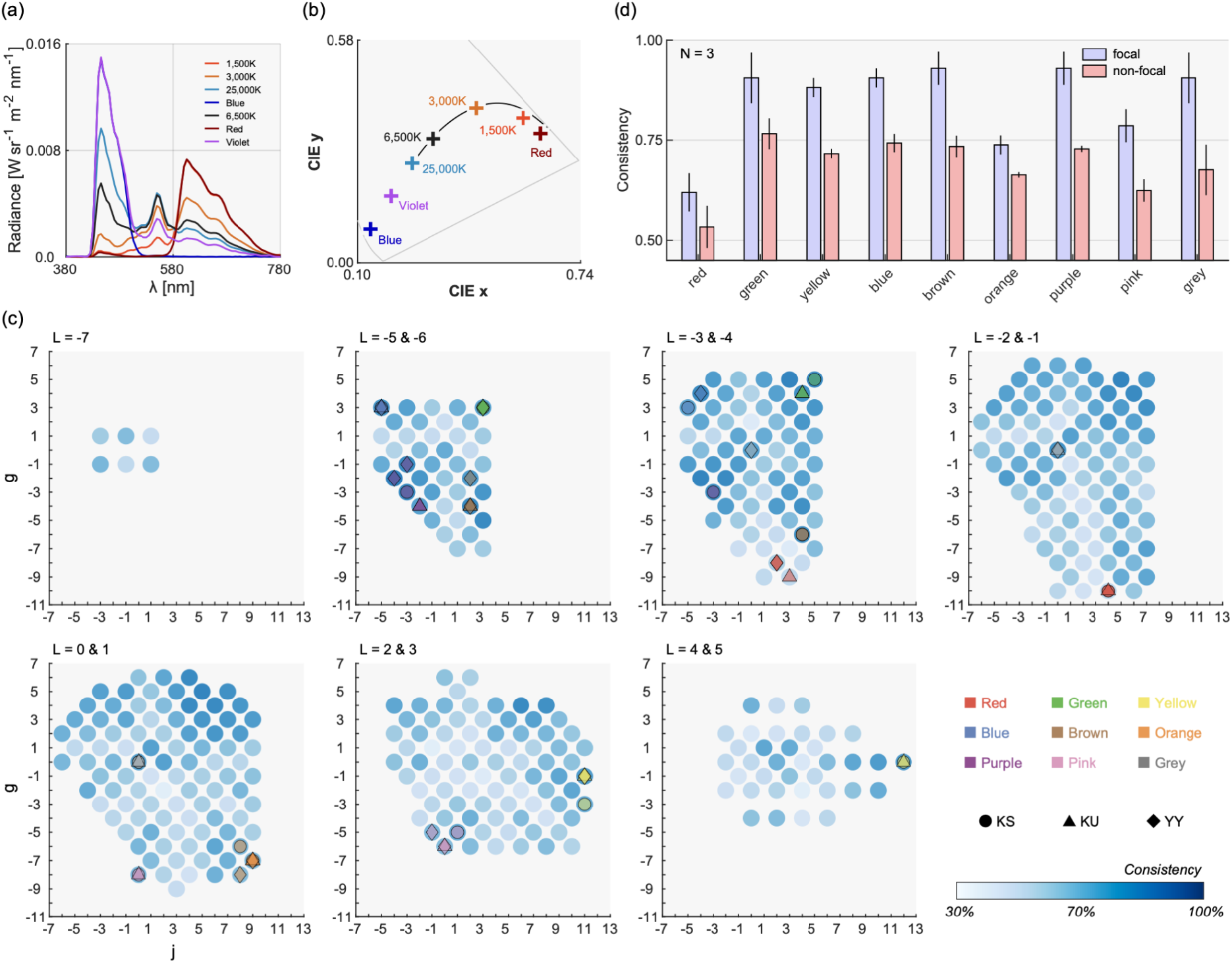
Experimental set-up and results in Experiment 1-2 where a large number of illuminants were used compared to Experiment 1-1. (a) Seven Illuminant spectra and (b) their chromaticity in CIE1964 xy chromaticity diagram. (c) The map of categorical consistency across illuminants for all color samples. Colored symbols indicate focal colors and shapes indicate different observers. (d) Consistency of color naming across illuminants for focal colors and non-focal colors. The error bars show ± 1.0 S.E across 3 observers.

### 4.2 Results and Discussion

Panel (c) in Figure 10 shows the consistency map across illuminant conditions for all color samples. For each color sample, there were 14 categorical responses (7 illuminants × 2 sessions) from which we calculate the consistency of color categories. The patterns are similar to Experiment 1-1 as shown in Figure 8, but the trend that focal colors coincide with highly consistent regions seems clearer. Panel (d) shows the averaged consistency for focal colors and non-focal colors, which again shows very similar trends to Experiment 1 (Figure 9 (a)). A two-way repeated-measures ANOVA confirmed that there is a significant main effect of focality (*F*(1, 2) = 100.4, *p* = 0.00981) and color category (*F*(8, 16) = 9.40, *p* = 0.000089). The interaction between the two factors was not significant (*F*(8, 16) = 1.01, *p* = 0.466), meaning that focal color had higher consistency than non-focal color for all color categories. To summarize, this additional experiment consolidated the main finding in Experiment 1-1.

## 5. Experiment 2: Asymmetric color matching

One major finding in Experiment 1 was that categorical color constancy holds remarkably well, especially under *3,000K* and *25,000K*. However, there are at least two potential reasons for this. First, it is possible that the color appearance of color samples did not change much under test illuminants compared to the *6,500K* condition, which consequently induces little categorical change. This first possibility shows that it is the appearance-based color constancy that held well, and the invariance of color category is simply a by-product. A second possibility, which we think is more interesting, is that though color appearance shifted influenced by the illuminant change, color categories did not change as they are inherently more tolerant than color appearance. This second possibility highlights the advantage to use a color category in color constancy mechanisms. Thus, to contrast the difference between appearance-based color constancy and categorical color constancy, Experiment 2 directly measured how much color appearance changed under a test illuminant using an asymmetric color matching task well-established in the field (e.g. Brainard & Wandell, 1992).

### 5.1 Conditions and Procedure

Figure 11 shows the scene configuration in Experiment 2. The scene consists of two parts. On the left-side, a test color sample was lit by a whole illumination. On the right-side, a reference color sample (*L, j, g* = 0) was lit by a spot illumination. For a surrounding region on both sides, we used the same gray color paper as Experiments 1. Observers viewed the scene with their forehead attached tightly to the black partition wall so that the right eye sees only the matching scene and that the left eye sees only the test scene (dichoptic presentation). For the test whole illumination, we used the five illuminants used in Experiment 1-1, namely *6,500K, 3,000K, 25,000K, Red* and *Blue* illuminants. For the test color sample, we used color samples named gray under each test illuminant, which allowed us to measure how the appearance of categorical achromatic points shifted depending on the illuminant color. The gray color samples were located on L = 0 and 1 planes for KU (51 samples in total across 5 test illuminants) and L = -1, 0 and 1 planes for YY (61 samples) and KS (31 samples). The observer’s task was to change the color of matching spot illumination by adjusting the intensity of R, G and B phosphors of the projector independently until the appearance of the reference and test color sample matched. Two matchings per color sample were obtained. There was no time limitation for each trial.

**Figure 11:**
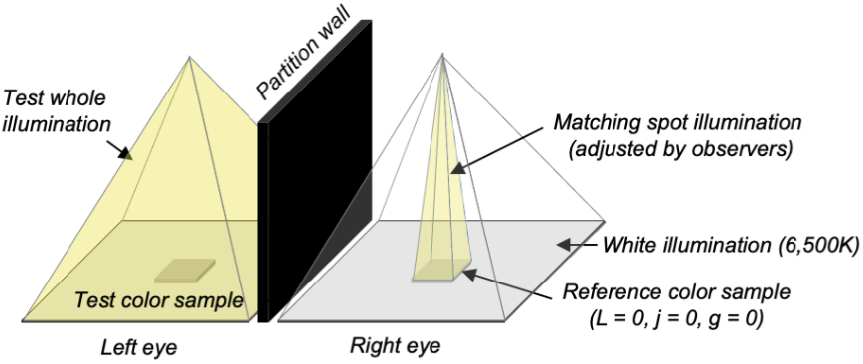
Experimental setup for asymmetric color matching. The left side is a test scene where the test color sample was illuminated by one of 5 illuminants used in Experiment 1 (*6,500K, 3,000K, 25,000K, Red* and *Blue*). For the test color sample, we used a color sample named gray under each test color illuminant. The right hand had a reference color sample (*L* = *j* = *g* = 0) that was illuminated by the spot illuminant, whose R, G and B values were independently adjusted by observers until the appearance of test and reference color samples matched.

### 5.2 Results

Figure 12 (a) - (c) shows matching results for each observer. Each colored small circle depicts the averaged matching point across two repetitions and diamonds show mean chromaticity across all settings for each illuminant. The ellipse was fitted to cover the data points in each illuminant condition with 95 % confidence intervals. For (a) observer KS, the matching points were clustered closely across all illuminant conditions. In contrast, for (b) KU and (c) YY, matching points in each illuminant condition were separated from each other. We quantified this separation using *DE2000* between *6,500K* (grey diamond) and other illuminants (colored diamonds) as shown in panel (d). Each bar shows an average across three observers. A one-way repeated-measures ANOVA showed that the main effect of illuminant color is significant (F(3,6) = 6.48, p = 0.0260), but multiple comparisons confirmed that there is no significant difference for any pair (Bonferroni’s correction). The mean *DE2000* value across participants was at least 6.21 (*3,000K* condition) and around 17 for *Red* and *Blue* illuminant conditions. These results show that shifts of color appearance are 17 times larger than one *jnd*, meaning that the appearance of the color samples named gray under Red and Blue illuminants are visibly different from the appearance of color samples named gray under 6,500K. Despite such large appearance differences, those color samples were still all named gray. To sum, these results suggest a category-based constancy is substantially more robust than appearance-based constancy. This greater robustness of categorical color constancy occurs potentially because categorical color naming is more tied to the judgment about the inherent property of an object as opposed to the color appearance of the object (so-called hue and saturation match vs. paper match, see Reeves et al., 2008 for further discussion).

**Figure 12:**
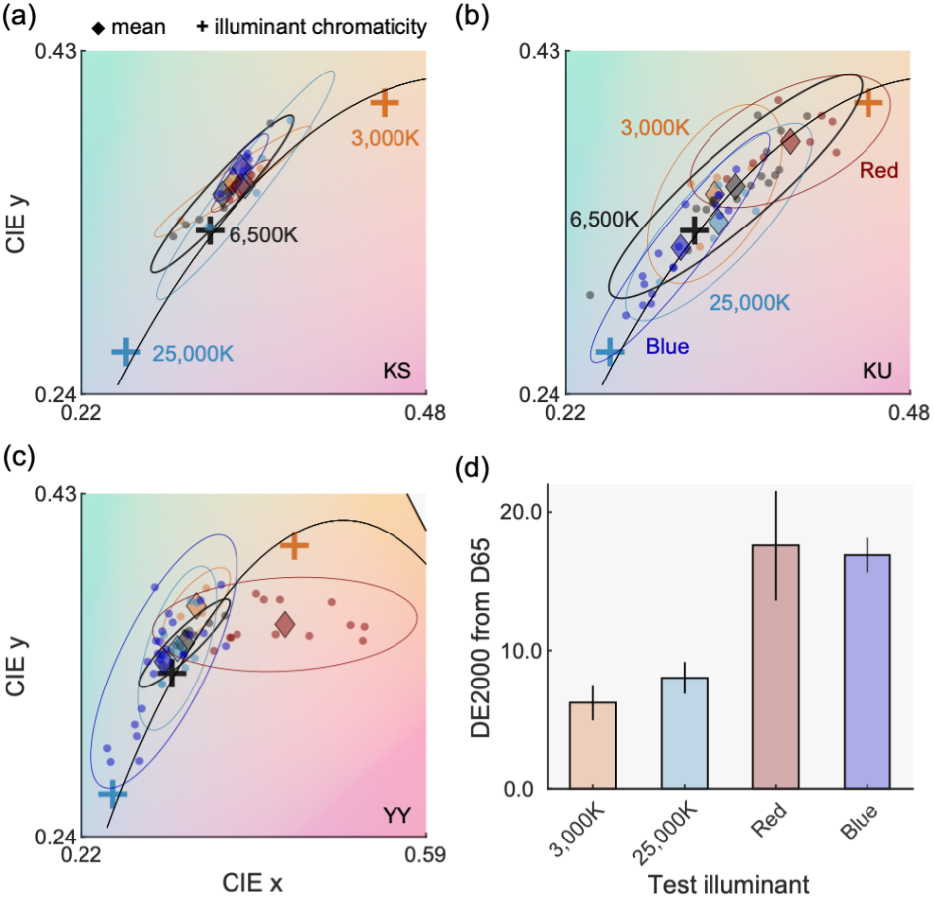
Results of asymmetric matching for (a) observer KS, (b) KU and (c) YY. Each small circle shows one matching. The diamond symbols depict the average across all matchings. Ellipses were fitted to data points with 95% confidence intervals. Notice that the x axis range is wider for (c) than (a) and (b). (d) DE2000 between mean matching points for 6,500K condition (gray diamond in panels (a) - (c)) and other illuminant conditions (colored diamonds). Each bar shows an average across three observers and error bars show ± 1.0 S.E across observers.

## 6. General Discussion

The present study measured categorical color naming patterns using a fairly large number of color samples placed under an extensive set of illuminants which allowed us to comprehensively quantify the degree of categorical color constancy. A spot illuminant methodology was introduced and shown to be effective to measure categorical patterns under different illuminants while silencing color constancy mechanisms. We further conducted an asymmetric matching experiment to measure how much color appearance shifted within the color samples that were all named gray. We found that categorical color constancy works generally well, and invariant categorical regions curiously overlap with focal color regions.

These results are largely in line with past studies (Olkkonenn, Hansen & Gegenfurtner, 2009; Olkkonnen et al. 2010). Furthermore, we found that focal colors do not necessarily locate around the centroid of color category, suggesting that focal color is stable for a nontrivial reason. Instead, focal colors may serve as a categorical anchor in mechanisms that mediate categorical color constancy. Results in the asymmetric matching experiment revealed a surprising amount of variation in color appearance among colors that were named gray, especially in illuminant conditions that are outside of the range of natural illuminant variation. The knowledge as to which regions in a color space remain categorically invariant is valuable in basic research, but it could be also used to inform ergonomic industrial designs. For example, color designers can pick a color from the invariant regions in the color space so that the color category of products stays the same under different lighting conditions.

While we generally observed quite a high degree of categorical color constancy, constancy indices shown in Figure 6 were still substantially lower than the upper bounds. One potential reason might be that there was not enough information about illuminant influence in the scene. Whilst the gray surrounding area would provide a direct cue to the color of illuminant, having some color variations in surrounding context also might have improved color constancy (Maloney & Wandell, 1986; Morimoto 2021). Another candidate would be the color of test illuminants we sampled. It is worth noting that in a previous study where temporal daylight variation within a day was measured (Morimoto et al, 2022), we found that most daylights fell within the range between *4,500K* and *20,000K*. Thus, it is possible that the natural illuminants we used (*3,000K* or *25,000K*) may have induced too large of illuminant changes. In retrospect, the use of less extreme correlated color temperature might have increased the degree of color constancy. However, conversely it may be surprising that a relatively high degree of color constancy was found even under these large illuminant changes.

We here list some limitations in the present study for future studies. Firstly, *OSA* color samples were used in all experiments. We chose the system mainly because of their perceptual uniformity, but categorical naming patterns are inherently constrained by the geometry of the color space and we indeed observed that the number of responses differed from one color category to another as shown in Figure 4 (b) and (c). This observation suggests that the use of finer scale might be desirable around a category where smaller numbers of responses were recorded (e.g. red or yellow). Consequently, we should not put too much emphasis when interpreting results for these categories. For example, the focal colors were found to be close to the categorical centroids for small categories (Figure 9 (b)), but this is more likely to stem from the property of stimuli than the nature of focal colors. Past studies also have emphasized the importance of controlling low-level visual information when measuring categorical influences (see Witzel & Gegenfurtner 2011; Witzel & Gegenfurtner 2015). Ultimately, the stimulus dependency of our findings is not exactly clear until other surface sets are tested, but it should be noted that Boynton et al. (1989) reported that color naming patterns did not differ in a significant way when Munsell color chips were used instead.

Another limitation regarding stimuli is that illuminants we used were sampled from the black-body locus and its extension. Though Olkkonenn et al. (2009) and Olkkonnen et al. (2010) found little differences between blue-yellow illuminants and red-green illuminants, it is an interesting question whether our findings hold for illuminants sampled at different regions in a color space. Thirdly, the number of observers recruited in this study is relatively limited. This is because we used a large number of color samples and illuminant conditions, and we aimed to collect a large number of responses per observer (in total 8,480 responses). However, consequently the present study does not directly address the individual differences in color categories, which is currently a missing piece in the field of categorical color perception to our knowledge. Finally, the ethnicity of observers in this study is limited to Japanese, and it is an open question how much findings translate to other observers who have different linguistic origins.

As we have found significance of focal colors, their potential roles in our categorical color system have been the center of discussion in the field. Several unique features of focal colors have been discussed and in some cases questioned. Are focal colors consistent across languages (MacLaury 1997; Abbot, Griffiths, & Regier, 2016; Regier, Kay & Cook, 2005; Kuehni, 2007)? Do focal colors appear more salient (Regier et al. 2007; Witzel & Franklin 2014)? Are there similarities between unique hues and focal colors (Kuehni, 2001; Miyahara, 2003)? Moreover, in agreement with our findings, some studies reported the particularly high categorical constancy for focal colors (Olkkonenn, Hansen & Gegenfurtner, 2009; Olkkonnen et al. 2010) while others found no evidence for this effect (Witzel et al. 2016). In communication using color terms, focal colors may be particularly critical because they are the colors that people are likely to imagine first in their mind, for example when they are requested to pass a red coat. Thus, for efficient communication it is beneficial to have higher stability at focal colors. The final remaining open question is why focal color is stable. We confirmed that it is not simply because they are at the categorical center. Here we propose an alternative view that a categorically invariant region might have evolved as a focal region. In other words, our visual system has chosen a region in a color space that is invariant under illuminant changes as a focal color, to facilitate robust communication with others using color terms in highly mutable environments we live in. If we instead have focal colors highly variable across illuminant changes, we may not have been able to rely on our color naming system. Until more comprehensive tests are conducted, this should stay a hypothesis. Nevertheless this might shed light on the origin of focal colors as a perceptual anchor.

## Acknowledgement

We thank Kenji Yokoi for his help to run experiments. This work was supported by JSPS KAKENHI Grant Number 19K22881. TM is supported by a Sir Henry Wellcome Postdoctoral Fellowship and a Junior Research Fellowship from Pembroke College, University of Oxford. This research was funded in whole, or in part, by the Wellcome Trust (218657/Z/19/Z). For the purpose of open access, the author has applied a CC BY public copyright license to any Author Accepted Manuscript version arising from this submission.

## Data access

The raw experimental data will be available in a data repository at the time of publication.

## Reference

Abbott, J. T., Griffiths, T. L., & Regier, T. (2016). Focal colors across languages are representative members of color categories. Proceedings of the National Academy of Sciences of the United States of America, 113(40), 11178–11183.

Arend, L., & Reeves, A. (1986). Simultaneous color constancy. Journal of the Optical Society of America A, 3(10), 1743–1751.

Buck, S. L., Shelton, A., Stoehr, B., Hadyanto V., Tang, M., Morimoto, T., and DeLawyer T. (2016) Influence of surround proximity on induction of brown and darkness. Journal of the Optical Society of America A, 33, 12–21.

Berlin, B., Kay, P. (1969). Basic Color Terms: Their Universality and Evolution. University of California Press, Berkeley/Los Angeles.

Bornstein, M. H., & Korda, N. O. (1984). Discrimination and matching within and between hues measured by reaction times: some implications for categorical perception and levels of information processing. Psychological Research, 46(3), 207–222.

Boynton, R. M., & Olson, C. X. (1987). Locating basic colors in the OSA space. Color Research & Application, 12(2), 94–105.

Boynton, R. M., Maclaury, R. E., & Uchikawa, K. (1989). Centroids of color categories compared by two methods. Color Research & Application, 14(1), 6–15.

Boynton, R. M., Fargo, L., Olson, C. X., & Smallman, H. S. (1989). Category effects in color memory. Color Research & Application, 14(5), 229–234.

Brainard, D. H., & Wandell, B.A. (1992). Asymmetric color matching: how color appearance depends on the illuminant. J Opt Soc Am A., 9(9), 1433–48.

JP de Vries, Akbarinia, A., Flachot, A., & Gegenfurtner K. R. (2021) Emergent Color Categorization in a Neural Network trained for Object Recognition. bioRxiv, https://doi.org/10.1101/2021.06.28.450097

Drivonikou, G. V., Kay, P., Regier, T., Ivry, R. B., Gilbert, A. L., Franklin, A. & Davies, I. R. L. (2007), Further evidence that Whorfian effects are stronger in the right visual field than the left, Proceedings of the National Academy of Sciences of the United States of America, 104, 3, 1097–1102.

Foster, D. H. (2011). Color constancy. Vision Research, 51(7), 674–700.

Granzier, J. J. M., Brenner, E., & Smeets, J. B. J. (2009). Reliable identification by color under natural conditions. Journal of Vision, 9(1), 1–8.

Gilbert, A. L., Regier, T., Kay, P., & Ivry, R. B. (2006). Whorf hypothesis is supported in the right visual field but not the left, Proceedings of the National Academy of Sciences of the United States of America. 103, 2, 489–494.

Gilbert, A., Regier, T., Kay, P. & Ivry, R. (2008), Support for lateralization of the Whorf effect beyond the realm of color discrimination, Brain and Language, 105, 2, 91–98.

Hansen, T., Walter, S., & Gegenfurtner, K. R. (2007). Effects of spatial and temporal context on color categories and color constancy. Journal of Vision, 7(4), 1–15.

Kay, P., & Kempton, W. (1984). What Is the Sapir-Whorf Hypothesis? American Anthropologist, 86(1), 65–79.

Kay, P., & Regier, T. (2003). Resolving the question of color naming universals. Proceedings of the National Academy of Sciences of the UnitedStates of America, 100(15), 9085–9089.

Kay P., Cook R.S. (2016). World Color Survey. In: Luo M.R. (eds) Encyclopedia of Color Science and Technology. Springer, New York, NY. https://doi.org/10.1007/978-1-4419-8071-7_113

Kay, P., & Regier, T. (2007). Color naming universals: The case of Berinmo. Cognition, 102(2), 289–298.

Kuehni, R.G. (2001). Focal colors and unique hues. Color Research & Application, 26(2), 171–172.

Kuehni, R.G. (2007). Nature and Culture: An Analysis of Individual Focal Color Choices in World Color Survey Languages. Journal of Cognition and Culture 7 (3-4), 151–172.

Lindsey, D. T. & Brown A. M. (2006). Universality of color names. Proceedings of the National Academy of Science of America, 103, 16608–16613.

Lindsey, D. T. & Brown, A. M. (2009). World Color Survey color naming reveals universal motifs and their within-language diversity. Proceedings of the National Academy of Sciences of America, 106, 19785–19790.

Luo, M. R., & Rigg, G. C. B. (2001). The Development of the CIE 2000 Colour-Difference Formula: CIEDE2000. Color Research and Application, 26, 340–350.

Ma, R., Liao, N., Yan, P., & Shinomori, K. (2018). Categorical color constancy under RGB-LED light sources. Color Research and Application, 43(5), 655–674.

MacAdam, D. L. (1974). Uniform color scales. J. Opt. Soc. Am. 64, 1691–1702.

MacAdam, D. L. (1978). Colorimetric data for samples of OSA uniform color scales. J. Opt. Soc. Am. 68, 121–130

MacLaury, R.E. (1997). Ethnographic evidence of unique hues and elemental colors. Behavioral and Brain Sciences 20 (2): 202–203.

Maloney, L. T., & Wandell, B. A. (1986). Color constancy: A method for recovering surface spectral reflectance. Journal of the Optical Society of America A, 3, 29–33.

Miyahara E. (2003). Focal colors and unique hues. Perceptual and motor skills, 97(3 Pt 2), 1038–1042.

Morimoto T., Kusuyama, T., Fukuda, K. & Uchikawa, K. (2021). Human color constancy based on the geometry of color distributions. Journal of Vision, 21(3), 7, 1-28.

Morimoto T., Zhang C., Fukuda K., & Uchikawa K. (2022) Spectral measurement of daylights and surface properties of natural objects in Japan. Optics Express, 30(3), 3183–3183.

Olkkonen, M., Hansen, T., & Gegenfurtner, K. R. (2009). Categorical color constancy for simulated surfaces. Journal of Vision, 9(12), 1–18.

Olkkonen, M., Witzel, C., Hansen, T., & Gegenfurtner, K. R. (2010). Categorical color constancy for real surfaces. Journal of Vision, 10(9), 1–22.

Okazawa G., Koida, K., & Komatsu, H. (2011). Categorical properties of the color term “GOLD”. Journal of Vision, 11(8), 4.

Parraga C.A., & Akbarinia A. (2020). Color Name Applications in Computer Vision. In: Shamey R. (eds) Encyclopedia of Color Science and Technology. Springer, Berlin, Heidelberg. ttps://doi.org/10.1007/978-3-642-27851-8_404-1h

Reeves, A.J., Amano, K. & Foster, D.H. (2008). Color constancy: Phenomenal or projective?. Perception & Psychophysics 70, 219–228.

Regier, T., Kay, P., & Cook, R. S. (2005). Focal colors are universal after all. Proceedings of the National Academy of Sciences of the United States of America, 102(23), 8386–8391.

Regier, T., P. Kay, & N. Khetarpal. (2007). Color naming reflects optimal partitions of color space. Proceedings of the National Academy of Sciences USA 104 (4): 1436–1441.

Siuda-Krzywicka, K., Witzel, C., Chabani, E., Taga, M., Coste, C., Cools, N., … Bartolomeo, P. (2019). Color Categorization Independent of Color Naming. Cell Reports, 28(10), 2471-2479.e5.

Troost, J. M., & De Weert, C. M. M. (1991). Naming versus matching in color constancy. Perception & Psychophysics, 50(6), 591–602.

Uchikawa, K., Uchikawa, H., & Boynton, R. M. (1989). Partial color constancy of isolated surface colors examined by a color-naming method. Perception, 18(1), 83–91.

Uchikawa, K., & Boynton, R. M. (1987). Categorical color perception of Japanese observers: Comparison with that of Americans. Vision Research, 27(10), 1825–1833.

Uchikawa, H., Uchikawa, K., & Boynton, R. M. (1989). Influence of achromatic surrounds on categorical perception of surface colors. Vision Research, 29(7), 881–890.

Uchikawa, K., Emori, Y., Toyooka, T., & Yokoi, K. (2002). Color constancy in categorical color appearance [Abstract]. Journal of Vision, 2(7):548, 548a, http://journalofvision.org/2/7/548/, doi:10.1167/2.7.548.

Uchikawa, K., Yokoi, K., & Yamauchi, Y. (2004). Categorical color constancy is more tolerant than apparent color constancy [Abstract]. Journal of Vision, 4(8):327, 327a, http://journalofvision.org/4/8/327/, doi:10.1167/4.8.327.

Winawer, J., Witthoft, N., Frank, M., Wulund, L., Wade, A. & Boroditsky, L. (2007). The Russian Blues Reveal Effects of Language on Color Discrimination. Proceedings of the National Academy of Sciences of the United States of America, 104, 7780–5.

Witzel, C., & Gegenfurtner, K. R. (2011). Is there a lateralized category effect for color? Journal of Vision, 11(12), 16.

Witzel, C. (2019). Misconceptions About Colour Categories. Review of Philosophy and Psychology Vol. 10. Review of Philosophy and Psychology.

Witzel, C., & Franklin, A. (2014). Do focal colors look particularly “colorful”? Journal of the Optical Society of America A, 31(4), A365.

Witzel, C., & Gegenfurtner, K. R. (2015). Categorical facilitation with equally discriminable colors. Journal of Vision, 15(8), 1–33.

Witzel, C., van Alphen, C., Godau, C., & O’Regan, C. K. (2016). Uncertainty of sensory signal explains variation of color constancy. Journal of Vision, 16(15):8, 1–24.

Witzel, C., & Gegenfurtner, K. R. (2018). Color perception: Objects, constancy, and categories. Annual Review of Vision Science, 4, 475–499.

